# The role of gene expression in the recent evolution of resistance in a model host parasite system

**DOI:** 10.1101/102301

**Authors:** Brian K. Lohman, Natalie Steinel, Jesse N. Weber, Daniel I. Bolnick

## Abstract

Damage by parasites is a perpetual challenge for hosts, often leading to the evolution of elaborate mechanisms of avoidance, immunity, or tolerance. Host resistance can evolve via changes in immune protein coding and/or expression. Heritable population differences in gene expression following infection can reveal mechanisms of immune evolution. We compared gene expression in infected and uninfected threespine stickleback (*Gasterosteus aculeatus*) from two natural populations that differ in their resistance to a native cestode parasite, *Schistocephalus solidus*. Genes in both the innate and adaptive immune system were differentially expressed as a function of host population, infection status, and their interaction. These genes were enriched for loci controlling immune functions that we independently verified differ between host populations, or in response to infection. For instance, populations differ strongly in reactive oxygen (ROS) production, and we observed corresponding differences in expression of ROS-affecting loci. Differentially expressed genes also were involved in fibroblast activation, B-cell activation, and leukocyte trafficking. Coexpression network analysis identified two distinct immune processes contributing to stickleback resistance; several modules of genes are correlated with parasite survival while a different set of modules are correlated with suppression of cestode growth. Comparison of networks between populations showed resistant fish have a dynamic expression profile while susceptible fish are static. In summary, recent evolutionary divergence between two vertebrate populations has generated population-specific gene expression responses to parasite infection, which reveal a few immune modules likely to separately affect cestode establishment, and growth.

## Introduction

Helminths are a diverse group of parasitic worms, which often establish long lasting infections in their vertebrate hosts (Maizels, et al. 2004), despite host immune activity. Curiously, in many host-parasite systems, helminths can persist in some host genotypes, whereas other hosts successfully eliminate infections. Therefore, a key question in biology is, why does parasite resistance differ among host individuals or populations? Host resistance depends on a complex signaling cascade, starting with the detection of pathogen molecules or pathogen induced damage to host tissues, followed by activation of a diverse suite of innate and adaptive immune cell populations. These cells may proliferate, migrate, or produce molecules that signal to other immune cells or directly attack the parasite. If the infection is cleared, the host must down-regulate this costly response (Maizels and Yazdanbakhsh 2003; Maizels, et al. 2004; Anthony, et al. 2007; Gause, et al. 2013). Natural genetic variation in host resistance could arise from any of these diverse stages of an immune cascade.

Classically, the search for genes important to host immunity has been conducted in the lab using a combination of forward genetic experiments and screens for abnormal phenotypes (Beutler, et al. 2006; Beutler, et al. 2007). Such approaches typically identify genes in which natural or induced mutations lead to loss of immunological function. In contrast, natural selection provides a powerful genetic screen for alleles that confer adaptive benefits within the complex ecological milieu in which wild vertebrates evolved and currently live, including diverse stresses and coinfections (Beraldi, et al. 2007; Schielzeth and Husby 2014). Isolated host populations are often exposed to distinct local parasite species or genotypes, and consequently evolve divergent immune traits. Spatially varying coevolution thus leads to adaptive geographic variation in host genotypes and corresponding immune traits (Eizaguirre, et al. 2012; Stutz and Bolnick 2016) In contrast to lab knock-out screens, this natural genetic variation is more likely to entail genes whose alleles confer a change or gain of immune function. Loss of function is of course also a possibility, if parasites exploit a given host trait, or if a trait confers insufficient benefits to warrant its costs. By identifying these evolutionarily labile genes, biologists seek to understand the genetic and immunological mechanisms of vertebrate resistance to, and coevolution with, helminth parasites. The genes identified in this manner will be of interest not only for what they tell us about the basic biology of host parasite interactions, but also as a possible source of new therapeutic strategies for controlling parasitic infections or manipulating vertebrate immunity (Geary, et al. 1999; Geary, et al. 2015).

One way to identify genes favored by natural selection is to look for evolution of gene expression in response to infection. Recent advances in sequencing technology and genetic mapping have made this an accomplishable goal (Cookson, et al. 2009; Pasaniuc and Price 2016). Previous studies have uncovered variation in gene expression associated with disease in rat, mouse, and human populations (Hubner, et al. 2005; Barrett, et al. 2008; Emilsson, et al. 2008; Wijayawardena, et al. 2016), but few studies have used wild populations (Hawley and Altizer 2011; Pedersen and Babayan 2011; Viney, et al. 2015; Huang, et al. 2016). These studies are often underpowered, as the historically high cost of RNAseq library prep and sequencing limited biological replication (Todd, et al. 2016). Few studies of variation in disease in wild populations have included more than a single population (Pedersen and Babayan 2011) or considered the effect of exposure on those individuals who did not ultimately become infected. Finally, linking changes in gene expression to host immune function requires concurrent measurement of multiple immune phenotypes, which are also missing from the majority of existing studies of wild populations of hosts. Here, we seek to close this gap by testing the effect of exposure or infection on gene expression, using a large number of individuals from two populations with independent evidence of immune trait divergence.

We tested whether host genotype and infection status alter host gene expression, using the threespine stickleback fish (*Gasterosteus aculeatus*) and its native cestode parasite *Schistocephalus solidus* as a model host-parasite system. The cestode’s eggs are deposited into freshwater via bird feces, then hatch and are consumed by copepods, which are in turn consumed by stickleback. Cestodes mature only in stickleback peritoneum, their obligate host, then mate inside the gut of piscivorous birds. This life cycle can be recapitulated in the lab, permitting controlled genetic crossing (Schärer and Wedekind 1999) and controlled infections among host or parasite genotypes. There is naturally occurring variation in cestode infection rates among stickleback populations throughout their native range (MacColl 2009; Weber, et al. 2017). This is mirrored by differences in expression of a select few immune genes, between wild caught stickleback from six populations, and between wild caught fish with versus without cestodes (Lenz, Eizaguirre, Rotter, et al. 2013; Stutz, et al. 2015).

Recently, Weber *et al.* identified natural populations of stickleback with dramatically different resistance to *S. solidus*. Marine stickleback, which resemble the likely ancestral state for modern freshwater populations, rarely encounter the cestode because its eggs do not hatch in brackish water. These fish genotypes are therefore highly susceptible to infection in laboratory exposure trials. When marine stickleback colonized post-glacial freshwater lakes, they encountered cestodes and evolved increased resistance to infection by cestodes (Weber, et al. 2017).

However, not all derived freshwater populations are equally resistant. On Vancouver Island in British Columbia, Gosling Lake (Gos) stickleback are heavily infected by cestodes (50 to 80% of fish, per year, from 10 years of observations). In contrast, the cestode is absent in stickleback from nearby Roberts Lake (Rob) over the same period of time (18 km away) (Weber, et al. 2016). The first host (copepods) and terminal hosts (piscivorous birds, mostly loons and mergansers) are common in both lakes. Diet data from both lakes shows that Rob and Gos fish consume copepods at an equal rate (Snowberg, et al. 2015; Weber, et al. 2016). The difference in infection rates is therefore not likely to be merely ecological. Accordingly, (Weber, et al. 2017) used experimental infections to confirm that Rob fish are more resistant to infection than Gos fish. In the lab, cestodes infect Rob and Gos stickleback at statistically indistinguishable rates, but Rob fish greatly reduce cestode growth (by two orders of magnitude). Rob fish are able to subsequently kill established cestodes by initiating peritoneal fibrosis which sometimes leads to the formation of a cyst and cestode death. While the mechanism underlying this cestode growth suppression and killing is uncertain, potential correlates are suggestive. Lab-reared Rob fish (or, F1 hybrids with a Rob dam) have a higher granulocyte: lymphocyte ratio following infection. In Rob fish, a higher fraction of the granulocytes generate reactive oxygen species (ROS), and these constitutively produce more ROS than cells from Gos fish. ROS is thought to damage the cestode tegument, and ROS production was negatively correlated with cestode growth. This higher ROS production by Rob fish is constitutive rather than induced by infection.

Given the immune phenotypes that differ between Rob and Gos stickleback, we hypothesized that these populations would exhibit constitutive and infection-induced differences in gene expression. Furthermore, we expected these differences to involve differential expression of immune genes, particularly those involved in ROS production and fibrosis. To test these hypotheses, we quantified gene expression of head kidneys from lab-reared Rob and Gos stickleback from three treatments: control, exposed but uninfected, and infected by *S. solidus.* We tested for: i) genes whose expression differs constitutively between populations, ii) genes which are involved in general responses to cestodes shared by both host populations, and iii) genes whose expression depends on the interaction between host population and infection status. Genes whose expression depends on an interaction between population and infection status are prime candidates for explaining how these populations respond differently to cestodes, ultimately resulting in significantly different parasite burdens. Additionally, we tested for correlations between modules of coexpressed genes and immune phenotypes (e.g. ROS production and granulocyte: lymphocyte ratio). The number of correlated suites of genes and their correlations with various immune phenotypes can give insight into pathway level phenotypes for further study. In particular, we wish to know whether cestode establishment, and cestode growth, are correlated with similar or different gene expression modules implying a shared or separate immunological cause.

## Results and Discussion

We obtained mRNA from the head kidneys of stickleback from three experimental groups: unexposed controls (N = 16 and 19, Rob and Gos fish, respectively), exposed but ultimately uninfected stickleback (N = 21 and 16), or exposed and infected (N = 17 and 9) fish. Tissues from the latter two groups were harvested 42 days post exposure. See (Weber, et al. 2016) for full experimental methods. We focus on expression in head kidneys as it is the major site of immune cell differentiation in fish (Scharsack, et al. 2004; Fischer, et al. 2006; Scharsack, et al. 2007; Fischer, et al. 2013), and head kidney cell cultures were used to measure immune function independently of gene expression. Below we present and interpret candidate genes from our linear models (see Table 1 for summary statistics).

**Table 1.**
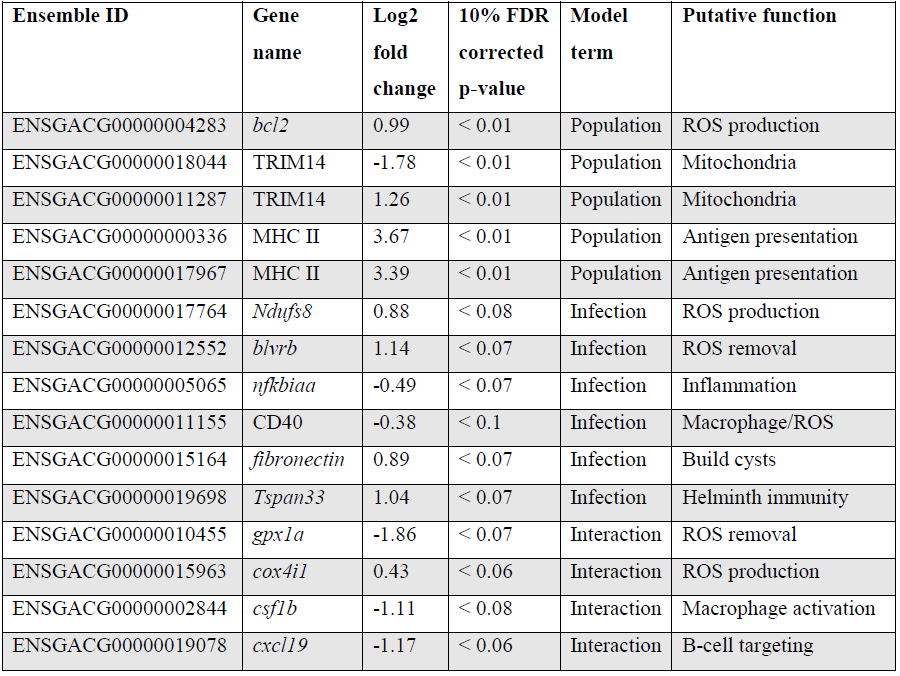
Candidate genes; Ensembl IDs, gene names, log2 fold changes, 10% FDR corrected p-values,term in the model for which they are significant, putative function, and interpretation

### Main effect of population on immune gene expression

Our negative binomial linear models (see Materials and Methods) identified 643 genes that were differentially expressed as a function of stickleback population (Wald, p < 0.1 after 10% FDR correction; 361 genes after 5% FDR). These main effects of population represent genes whose expression differs constitutively between populations (regardless of infection status). Because these differences occur in lab-raised fish, they represent heritable between-population differences in RNA abundance. Because we measured gene expression from the entire head kidney, expression differences could reflect evolved changes in gene regulation per cell, changes in cell population composition, or both). A caveat is that because we used first-generation lab-reared fish, we are as yet unable to rule out maternal or other epigenetic effects. However, comparison of Rob, Gos, and reciprocal F1 hybrids revealed little evidence for maternal effects on infection outcomes or immune traits (with the exception of granulocyte: lymphocyte ratio) (Weber, et al. 2016). So, we consider maternal effects unlikely for most of the differentially expressed genes documented here.

Previous studies have considered the effect of genotype on changes in stickleback immune gene expression in controlled lab infection experiments. However, these results are conflated with other factors such as environment (i.e., comparing wild-caught lake, stream, and estuary stickleback) and multiple exposures to parasites (Lenz, Eizaguirre, Rotter, et al. 2013). Host genotype was also considered in an experimental infection of honeybees, revealing significant host genotype effects on both gene expression and infection phenotypes (Barribeau, et al. 2014). Furthermore, host genotype effects could be potentially very important in mosquito-malaria interactions, including a unique example of dual-species trancriptomics (Choi, et al. 2014). Clearly host genotype effects in marcroparasite infection are worthy of future study.

Gene ontogeny (GO) showed that these differentially-expressed genes are significantly enriched for several categories related to mitochondrial respiration, which can affect ROS production (Figure 1, cellular components, Mann-Whitney U, p < 0.01 after 10% FDR correction). Rob lake fish also have higher expression of B-cell lymphoma 2 (*bcl2*, ENSGACG00000004283, log2foldchange = 0.99, Wald p 0.01 after 10% FDR correction), a mitochondrial membrane protein which mediates the release of ROS-producing cytochrome C into the cell and promotes cell survival in the presence of oxidative stress (Martindale and Holbrook 2002). We observed significant differences in expression of two copies of another mitochondrial adaptor, tripartite motif 14 (TRIM14). Surprisingly, expression of each gene copy changes in opposite direction between the two host populations (ENSGACG00000018044: log2fold change = −1.78, Wald p < 0.01 after 10% FDR correction, ENSGACG00000011287: log2fold change = 1.26, Wald p < 0.01 after 10% FDR correction). TRIM14 is part of the innate immune system (Zhou, et al. 2014), and has been shown to be under balancing selection among other populations of stickleback (Hohenlohe, et al. 2010). While the majority of differences in TRIM14 expression are constitutive population effects, there is a single copy depends on an interaction between population and infection status (see below). Together, the population differences in ROS-associated gene expression supports our observation of significantly greater ROS production in Rob stickleback. It is important to note that these genes are differentially expressed between populations regardless of infection status, consistent with prior observations that ROS production is constitutive, insensitive to infection status (Weber, et al. 2016).

**Figure 1.**
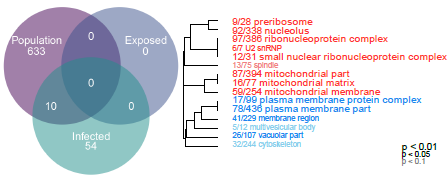
Linear modeling reveals differences between populations and by infection status (all genes p < 0.1 after 10% FDR correction). Genes which are differentially expressed between Rob and Gos are enriched for mitochondrial respiration (cellular components, Mann-Whitney U, p < 0.01 after 10% FDR correction). GO categories in red are upregulated, while blue indicates downregulated, relative to Gos. Numbers indicate genes present in category / total genes in category

Major histocompatibility complex II (MHC II) is a key element of the adaptive immune system, involved in pathogen recognition. Regardless of infection status, Rob fish have higher MHC II expression than do Gos fish, for two different copies of MHC II (ENSGACG00000000336: log2fold change = 3.67, Wald p 0.01 after 10% FDR correction, ENSGACG00000017967: log2fold change = 3.39, Wald p < 0.01 after 10% FDR correction). This difference in transcript abundance could be due to changes in the relative abundance of antigen presenting cells (APCs) such as macrophages, which express MHC II (Murphy 2011). To test this possibility we tested another statistical model to determine whether variance-stabilized expression of each MHC copy covaried with the proportion of granulocytes (as opposed to lymphocytes) in a head kidney primary cell culture, controlling for population and infection status. Rob fish have relatively more granulocytes when infected (Weber, et al. 2016), so we expected a positive correlation between MHC II expression and granulocyte production. Instead, the correlation was negative (ENSGACG00000000336: β = -0.0262, t = -1.76 ENSGACG00000017967: β = -0.022, t = -1.98). Our working model to explain this result is that Rob fish have constitutively higher abundance of MHC II in their head kidneys because they have higher numbers of antigen presenting cells regardless of infection status. When challenged by cestodes, Rob fish initiate a strong innate immune response, proliferating granulocytes but not antigen presenting cells. This infection-dependent proliferation of non-APC granulocytes may dilute the relative abundance of MHC transcript, resulting in the observed negative correlation between MHC and granulocyte abundance.

Previous work has focused on the role of MHC allelic variation in stickleback-parasite interaction, resistance, and local adaptation (Kurtz, et al. 2004; Wegner, et al. 2006; Lenz, Eizaguirre, Kalbe, et al. 2013). However, most of this work centers on MHC allelic composition and its correlation to infection and growth phenotypes (as a proxy for fitness). Many fewer studies quantify expression of MHC alleles. One study noted increased expression of MHC II in wild fish which were more heavily parasitized, especially when MHC allele diversity was low (Wegner, et al. 2006). However, only a single population of fish was considered, so our discovery of significant effect of population (Rob vs. Gos) on MHC II expression is therefore novel. It is important to note that stickleback have more than two copies of MHC AI. Previous sequencing efforts have suggested that stickleback have between 4 and 6 copies of MHC II (Sato, et al. 1998; Reusch, et al. 2004; Reusch and Langefors 2005). Because TagSeq does not sequence the entirety of the mRNA, we cannot distinguish all MHC haplotypes present in individual fish. It is therefore possible that differences in expression of the two variants described here is due to altered regulation of only particular alleles or paralogs.

### Transcriptomic response to exposure or infection

Surprisingly, no genes differed between control versus exposed-but-uninfected fish (Wald, p > 0.1 after 10% FDR correction). This could be because resistant fish quickly mounted a response to the cestode, eliminated the parasite, and then down-regulated immune function by our 42-day sample date. Or, early-acting resistance to the cestode may involve physical or chemical barriers to entry that entail constitutive gene expression or non-genetic effects (e.g., gut epithelial mucous, protective symbiotic bacteria, etc). Finally, early-stage infections may induce localized immune responses in the intestinal epithelium or peritoneum that are not reflected in head-kidney gene expression.

Once the cestode establishes in the peritoneum, however, it induces some shared changes in gene expression of all host genotypes. We identified 64 genes that were differentially expressed between control and infected fish (Wald, p < 0.1 after 10% FDR correction), across both host genotypes. Several of these genes are promising candidates because of their known role in host immunity. For example, infected fish increase expression of *ndufs8*, a component of complex I which is the main ROS producer in cells (ENSGACG00000017764: log2fold change = 0.88, Wald p < 0.08 after 10% FDR correction) (Procaccio, et al. 1997). Other subunits of complex I are more highly expressed in Rob fish regardless of infection status, consistent with their higher constitutive production of ROS. Therefore, *ndufs8* may be particularly important in regulating the induction of ROS in response to infection, because it is the only complex I subunit upregulated upon infection. ROS are reduced when they act on their targets, and the raw materials can be recycled through the biliverdin/billiruben redox cycle (Barañano, et al. 2002). Infected fish from both populations have higher levels of *blvrb* (biliverden reductase B), one of the two enzymes in this ROS-recycling system (ENSGACG00000012552: log2fold change = 1.14, Wald p < 0.07 after 10% FDR correction). This up-regulation should facilitate removal of ROS that may limit damage to host tissues, or facilitates subsequent ROS production.

Another important aspect of ROS-based immunity is the associated inflammation. Infected fish have decreased expression of *nfkbiaa* (nuclear factor kappa light polypeptide gene enhancer in B-cell inhibitor alpha a), which interacts with NF-kB to inflammation (ENSGACG00000005065: log2fold change = – 0.49, Wald p < 0.07 after 10% FDR correction). *Nfkbiaa* inhibits the pro-inflammatory NF-kB by either preventing NF-kB proteins from entering the nucleus, where they are active, or by blocking NF-kB transcription factor binding sites. NF-kB activation by TNFa or LPS reverses this binding and allows NF-kB to activate expression of pro-inflammatory genes (Verma, et al. 1995). Thus, decreased *nfkbiaa* expression suggests an increased inflammatory response following successful infection.

Infected fish also have a slight decrease in expression of CD40 (ENSGACG00000011155: log2foldchange = −0.38, Wald p < 0.1), a co-stimulatory molecule expressed on dendritic cells, macrophages, and B-cells, which activates T-and B-cells (Murphy 2011). Previous studies have suggested that helminths could potentially suppress stickleback adaptive immunity (Scharsack, et al. 2007), and the downregulation of CD40 is one plausible mechanism. Alternatively, fish with inherently lower CD40 expression may be more susceptible to infection. This raises a broader question that we are not yet able to answer, but which warrants further study: to what extent are the expression differences between infected and control fish a result of host immune response or parasite immune suppression?

CD40 expression is not limited to immune cells, but can also be expressed in fibroblasts (Grewal and Flavell 1998), so its precise function in stickleback infection by cestodes is unclear. This dual role is intriguing because fibroblast activation is associated with the formation of fibrotic cysts that encapsulate cestodes (Zeisberg, et al. 1999). These cysts likely restrict cestode movement and concentrate ROS while limiting damage to host tissues. Recall that this is a population specific defense, exhibited by Rob but not Gos fish (Weber, et al. 2016), and in this statistical contrast the effect of population is averaged. In addition to changes in CD40, our linear model identified an increase of expression of *fibronectin* in infected Rob fish, which contributes to fibrinogen production to build cysts (Gratchev, et al. 2001; Anthony, et al. 2007) (ENSGACG00000015164: log2fold change = 0.89, Wald p < 0.07 after 10% FDR correction).

Adaptive immune system genes also respond to cestode infection. *Tspan33* has recently been shown to be a marker for activated B-cells in vertebrates (Hevezi, et al. 2013; Perez-Martinez, et al. 2015). The presence of activated B-cells indicate the host immune system has recognized the parasite and is actively mounting a defense. In our study, infected fish show higher levels of *tspan33* compared to controls (ENSGACG00000019698: log2fold change = 1.04, Wald p < 0.07 after 10% FDR correction). Increased expression of *tspan33* in infected fish is consistent with increased activation of B-cells, an integral part of the adaptive immune response.

### Genotype by infection interactions

The higher resistance to *S. solidus* infection in Rob compared to Gos stickleback could be due to constitutive differences in gene expression (as documented above), or differences in the induced immune response to infection. The latter can be detectable via interactions between host genotype and infection status. Linear modeling results identified 16 genes significant for this interaction (Wald p < 0.1 after 10% FDR correction). Most of these genes are known to affect the immune traits that (Weber, et al. 2016) already showed are divergent between Rob and Gos fish. For example, glutathione peroxidase 1a (*gpx1a*) is an enzyme that degrades hydrogen peroxide, a type of ROS, into glutathione and water (Turrens 2003). Expression of *gpx1a* in Gos fish increases upon infection, and therefore should tend to decrease the amount of ROS (hydrogen peroxide) available to defend against cestodes (ENSGACG00000010455: log2foldchange = -1.86, Wald p < 0.07 after 10% FDR correction). We speculate that this proactive down-regulation upon infection might be a tolerance response to mitigate autoimmune damage by Gos fish, which are commonly infected and therefore might not be able to tolerate a strong ROS response. The cytochrome c complex produces ROS (Turrens 2003), and we see increased expression of Cytochrome c oxidase subunit IV (*cox4i1*) in Rob fish that are infected, while Gos fish decrease expression (ENSGACG00000015963: log2foldchange = 0.43, Wald p < 0.06 after 10% FDR correction). This gene expression data is consistent with our phenotypic data showing that Rob fish have more ROS producing macrophages than Gos fish, and more ROS per cell. This *cox4il* up-regulation in Rob fish may be amplified by population differences in *bcl2* (see above). Oddly, we do not observe a significant infection-induced increase in ROS production in fish of either genotype. This discrepancy may reflect our head-kidney cell-culture based ROS assay, which does not rule out changes in ROS *in vivo* or in other tissues. The one contrary result involves colony stimulating factor 1b (*csf1b)*, a paralog of *csf1/mcsf*, a well-studied regulator of monocytes in mammals (Akagawa, et al. 2006). *csf1* increases the production of head kidney leukocytes (which includes granulocytes) in trout (*Oncorhynchus mykiss*)(Wang, et al. 2008). In our study, *csf1b* is down regulated in infected Rob fish even though they have more granulocytes (which includes macrophages) relative to either Gos fish or to uninfected Rob fish (ENSGACG00000002844: log2foldchange = -1.11, Wald p < 0.08 after 10% FDR correction). This discrepancy may be resolved by recognizing that we examined a single time point post-exposure. It is likely that Rob fish initially increase *csf1b*or another gene to drive the granulocyte proliferation that we observe after infection. The down-regulation of *csf1b* 42 days after infection could be a homeostatic mechanism to suppress further macrophage proliferation, after they already reached sufficient abundance. Further time series analyses would be necessary to resolve this hypothesis.

Finally, adaptive immune system genes also exhibit population specific responses to infection. Activated B-cells are critical to mounting an adaptive immune response, and they are targeted by various cytokines (Murphy 2011). When challenged by cestodes, Rob fish increase expression of C–X–C motif chemokine ligand 19 (*cxcl19*). In contrast, cestode infection reduces *cxcl19* expression in Gos fish, which otherwise exhibit constitutively higher expression than Rob fish (ENSGACG00000019078: log2foldchange = -1.17, Wald p < 0.06 after 10% FDR correction; Fig 2). Ligands with this motif induce migration of leukocytes (Belperio, et al. 2000). Literature on *cxcl19* is rare, but it has been suggested that the zebrafish cxcl19 gene is orthologous to Il-8, a major mediator of leukocyte migration to sites of inflammation (Wang, et al. 2008). Regardless of whether *cxcl19* is involved specifically leukocyte trafficking to sites inflammation or increasing migration of leukocytes in the absence of inflammation, both of these immune mechanisms could play an important role in defense against cestodes.

**Figure 2.**
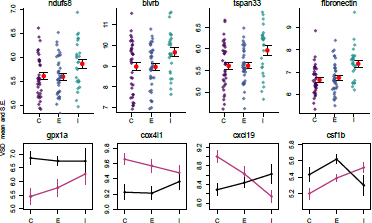
ROS production and B cells respond to infection by cestodes, both in a population independent,and dependent manner. Y axis is variance stabilized data, the product of log transforming and library size correcting raw gene counts. C = control, E = exposed, I = infected. Black lines are Rob and magenta lines are Gos.

### Expression – trait covariance

We tested for correlations between patterns of gene expression and immune/cestode phenotypes using weighted gene co-expression network analysis (WGCNA). WGCNA provides an unbiased data-driven hierarchical clustering of genes with similar expression patterns, thereby reducing the number of genes under consideration (reduced multiple test correction) and identifying functionally similar gene modules which can be used for further statistical analysis (Langfelder and Horvath 2008). WGCNA is a more appropriate analysis for incorporating additional immune phenotype data that was collected during the infection experiment not only because of its continuous nature (vs. the categorical predictors of population and infection status) but also because the correlation between suites of co-expressed genes and traits is estimated independently for each trait, rather than simultaneously (as under the linear modeling framework), resulting in lower unexplained variance to be assigned to other traits. We used a two-step process, first looking for general pathways and subsequent correlations to phenotypes by using all samples to construct a signed network. Second, we tested for genotype-dependent network structure and module-trait correlation by building signed networks for each population of stickleback. The latter case may be especially pertinent if the regulation of gene expression plays a strong role in the genotype dependent response to infection. To explore this, we calculated module similarity between the Rob and Gos signed networks as the fraction of genes shared between any two given modules.

When all samples were combined to build a single signed co expression network, WGCNA analysis revealed modules that were correlated with host population, ROS production, infection status, number of cestodes, total cestode mass, the density of cells in host head kidney, the frequency of granulocytes/lymphocytes, the fraction of cells gated into various subpopulations including precursors, myeloid, and eosinophils, and finally, host families (data not shown, but this highlights the need to include family as a nested factor in the linear modeling of DESeq2).

Population differences are mainly captured by the turquoise, green, magenta, and greenyellow modules, with lesser contributions by the blue and red modules (Figure 3). These population-dependent modules have connections to population-dependent phenotypes such as ROS production. For example, top kME genes in the turquoise modules (positive correlation with Rob) include ROS producing cytochrome c oxidase genes and ROS recycling *gpx1a* (Figure 2). Together, we would expect the action of these genes to increase ROS levels. As expected the turquoise module has a positive correlation with ROS production (r = 0.36 p = 2e–4, Figure 3).

**Figure 3.**
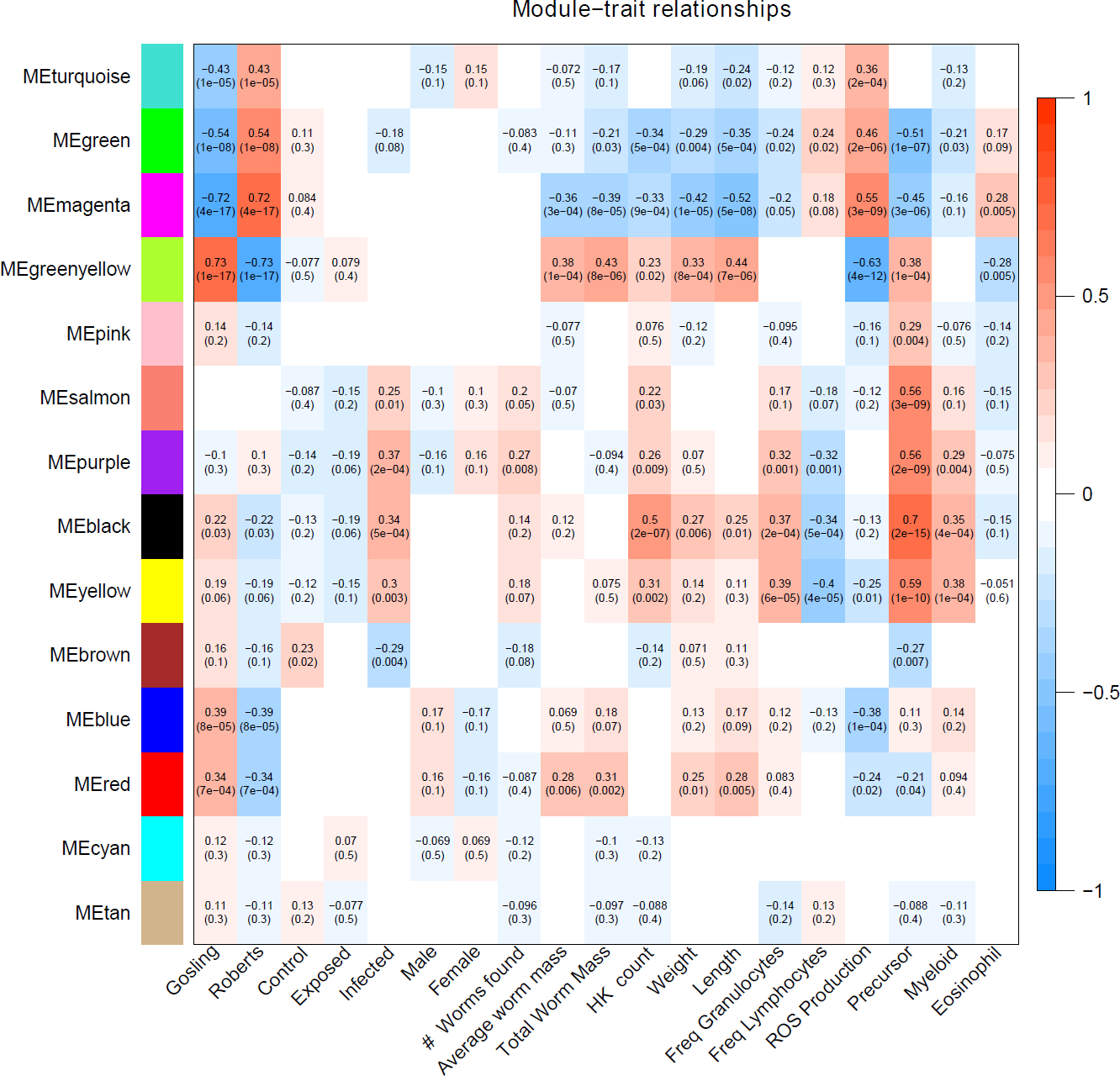
Weighed co expression gene network analysis suggest that host response to cestodes involves two traits, the initial immune response (salmon, purple, black, yellow, brown) and the control over parasite growth (magenta and greenyellow). Each cell indicates the correlation between the module and the p-value for that correlation (in parentheses). Correlations weaker than 0.07 were omitted.

Some module-trait correlations reinforce prior inferences about the immunological basis of stickleback resistance to the cestode. For example, the black module has modest to strong correlations with infection status, cell population phenotypes including density of all cells, precursors, and myeloids but is not correlated to ROS production (r = 0.13 p = 0.2, Figure 3). In contrast, the magenta and greenyellow modules are correlated with ROS production and also with cestode size, but much less so with cell population phenotypes and not at all with infection status (Figure 3). These observations imply that stickleback prevention of cestode establishment, and suppression of cestode growth, entail two distinct immune pathways (innate response and ROS production, respectively).

The magenta module has a modest, but strongly significant correlation to the fraction of cells that are eosinophils (r = 0.28, p = 0.005). A recent review highlighted the importance of eosinophils in host-helminth interactions. Specifically, at the site of host tissue damage, eosinophils are primed by fibronectin, and produce a variety of proteins which are toxic to helminths. Furthermore, the diversity of eosinophil cell surface receptors makes them central to mediate the inflammation response. Finally, helminths appear to have a number of anti-eosinophil proteins which both evade and dampen host response to the presence of helminths (Shin, et al. 2009).

Constructing separate signed coexpression networks for each population reveals dramatic differences in network structure. Using the same construction parameters for each population, the Rob network resembles the combined population network, showing strong module-trait correlations for worm size, cell population phenotypes, ROS production, and sex across many different modules. In stark contrast, the Gos network is much more static, with only a subset of traits which were previously significant correlated to a single module. Overall, Gos fish have many fewer total modules and correlations between modules and traits are much weaker. To estimate relationships between networks we calculated module similarity between every pair of Rob and Gos modules by shared gene membership. Broadly, large modules in Gos are split into many smaller modules in Rob. Thus, we conclude that there are two possible outcomes: 1) the Rob fish have evolved a more modular and dynamic repertoire of expression with which to fend off cestodes, or 2) the Gos fish evolved to maintain constitutive gene expression in response to cestodes, instead adopting a tolerance strategy (see supplementary materials for population specific module-trait heat maps and module similarity heat map).

Our WGCNA results confirm observed population differences in immune function and patterns of gene expression in the linear modeling. Furthermore they support our hypothesis that there are two traits involved in stickleback resistance to cestodes: 1) innate immune response to prevent cestode establishment and 2) limiting worm growth once cestodes become established. These two traits separate into distinct modules of gene expression, each enriched for genes with immunological function matching *a priori*expectations. This two trait perspective refines the question of variation in cestode prevalence among stickleback populations from by focusing attention on early and late stage infection. Future studies will be needed to describe the full time-series of gene expression as exposure and infection proceeds in these study populations, and to establish directionality of the interaction between cestodes and stickleback. While others have argued that cestodes are the primary drivers of coevolution (Scharsack, et al. 2007), only by sequencing both host and parasite mRNAs can we hope to detail this interaction at the molecular level. Host genotype by parasite genotype interactions offer a promising opportunity for further study. Such GxG interaction was recently described in the stickleback-cestode system, but only documented growth phenotypes, and did not attempt to describe the genetic basis for such traits (Kalbe, et al. 2016). We maintain that this type of study provides a means to identify evolutionarily labile genes which underlie beneficial shifts in immune function across the geographic mosaic of host-parasite coevolution.

### Summary

Using a large scale controlled laboratory infection experiment, we find changes in gene expression between two host populations, and as a function of infection status. For a smaller portion of genes, the expression response to infection differed between the two host populations. These findings are consistent with observations of host immune function in said infection experiment (Weber, et al. 2016). ROS production and recycling, B cell activation and targeting, and fibrosis appear to play important roles in stickleback defense against cestodes. Our analysis also suggests that host resistance involved two components; response to challenge by cestodes, and control over cestode growth. Furthermore, differences in co expression networks between populations suggest that either Rob fish have evolved a more elaborate expression profile or that Gos fish are shutting down expression to tolerate cestodes. Our results not only suggest a mechanistic link between host immune phenotypes and candidate genes, but also provide the foundation for studying the direct effects of host alleles on parasite fitness.

## Materials and Methods

### Sample collection, sequence library construction, and analysis of flow cytometry data

Stickleback samples were generated as part of a large lab infection experiment (Weber, et al. 2016). Briefly, head kidneys were dissected from stickleback and preserved in RNAlater at −20ºC. Libraries were constructed according to (Lohman, et al. 2016). Samples were sequenced on the Illumina HiSeq 2500 at the Genome Sequence and Analysis Facility at the University of Texas at Austin, producing ~6.7M raw reads per sample. See (Weber, et al. 2016) for ROS and flow cytometry methods. Flow cytometry data was analyzed using FlowJo software (Treestar). Granulocyte and lymphocyte populations were defined based on gating described in (Weber, et al. 2016). Precursor, myeloid, and eosinophil populations were defined using gating as described in (Wittamer, et al. 2011).

### Bioinformatics

TagSeq reads were processed according to the iRNAseq pipeline (Dixon, et al. 2015), using version 79 of the stickleback transcriptome from Ensemble, producing 9077 genes among all samples. Transcriptome annotations were based on the UniProtKB database (http://www.uniprot.org/help/uniprotkb) and followed previously described procedures (Dixon, et al. 2015). Code for the iRNAseq pipeline can be found here:https://github.com/z0on/tag–based_RNAseq. Code for the annotation pipeline can be found here:https://github.com/z0on/annotatingTranscriptomes.

### Statistical analysis with DESeq2

We scanned for outliers using arrayQualityMetrics (Kauffmann, et al. 2009) and removed 1 sample because of insufficient read depth. (final N = 98). To test for differential gene expression we used the following model in DESeq2:

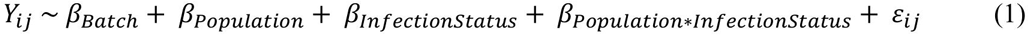

where *Y_ij_* is the count of gene *i* in individual *j*, *β_Population_* is a fixed effect with two levels: Rob and Gos, *β_InfectionStatus_* is a fixed effect with three levels: control, exposed (but not infected), and infected, and full-sibling families are nested within populations. β_Batch_ is the lane on which samples were sequenced. An additional predictor β_Sex_ was included for genes when appropriate (lower AIC score), and improved the model fit of 839 genes total. We fit the full model (including sex) to all genes and then extracted only the 839 that were improved by the addition of sex and looked for significant p values for main effects and interactions. With the full model, 67 of these ‘sex improved’ genes were significantly different between populations. No genes were significant for either exposure or infection, and 1 gene was significant for the interaction of population and infection status (myosin 5ab, ENSGACG00000006025: log2foldchange = 2.98, Wald p = 0.07). All p-values were multiple test corrected using 10% FDR. Although fish from the controlled infection experiment were exposed to three different parasite genotypes (each family exposed to only one parasite genotype), we are only interested in the host response to any parasite, and therefore average across parasite genotypes by simply not including this as a term in our linear model.

### Gene Ontogeny with GO_MWU

We used the Mann-Whitney U test for GO analysis. This approach has been described (Wright, et al. 2015) and the code for analysis can be found here:https://github.com/z0on/GO_MWU

### Weighted coexpression gene network analysis

Raw read counts were normalized using limma (Ritchie, et al. 2015) for input into WGCNA (Langfelder and Horvath 2008). Following the walkthrough in (Langfelder and Horvath 2008), we built a signed network with a soft thresholding power of 7, and a minimum module size of 30 genes. Following dynamic tree cut, we merged modules with greater than 80% similarity, producing 14 modules. We separated Rob and Gos samples and repeated this process with the same parameters.

### Caveats and limitations

Our study flips the traditional search for immune candidate genes from inbred lab strains to wild populations, using historical natural selection as a tool to screen for changes in gene expression associated with parasite infection. While our host-parasite model system is powerful, it does have some limitations. The reference genome is of generally good quality but annotation is lacking (approximately 22.5% of the entries in the stickleback genome are either unnamed or labeled as novel genes). Thus, GO analysis is performed after assigning GO accession terms by BLAST homology, rather than functional verification, a common solution for non-model systems. The features of the stickleback genome may be missing potentially interesting immunological genes which are sufficiently diverged from human or mouse genes and therefore may be unannotated. In particular, the number and location of MHC II paralogs remains uncertain, illustrating need for genome sequence improvement.

Our linear modeling with DESeq2 employs appropriate FDR correction, but we choose to accept higher than ‘standard’ p-values associated with LFC because of the direct connection between candidate genes and independently observed immune phenotypes. If, for example, we had not measured immune phenotypes, we would not accept log2 fold changes in expression with associated p-values greater than 0.05 but less than 0.1. Furthermore, we chose a very low base min mean filter because we have high confidence in detecting lowly expressed genes (Lohman, et al. 2016). We also wished to include more genes in our enrichment analysis and this also detracts from our power due to multiple test correction. We accordingly accept slightly larger than normally allowed p-values. Our TagSeq based approach has been shown to be at least as good as total RNAseq methods (having an equal or higher correlation between observed and known values of a spike in control), but does not account for splice variants or copy number variation, which may be potentially important in the evolution of immune responses.

Our study used tissue from a single organ (head kidneys) for both gene expression and immune phenotype measures. Head kidneys are a crucial hematopoetic organ in fish, but analysis of other tissues may produce different results. Moreover, head kidneys contain multiple immune cell populations that we are unable to sort effectively for cell-type-specific expression studies. We do use cell population counts (proportion granulocytes versus lymphocytes) as a covariate, which as noted for MHC weakly contributes to expression variation of a few genes. But, lacking monoclonal antibodies to many immune cell receptors in stickleback, we cannot readily distinguish among finer subdivisions of cell types. This resource limitation, typical of most non-model organisms, limits our ability to statically detect effects of cell population composition on expression.

## 4 Supplementary Material and Data to be Deposited

1. Fish meta data
2. Excel spreadsheet containing all DEGs
3. Input data for linear modeling (counts and design matrix)
4. Code for raw reads processing (generating gene counts)
5. Code for linear modeling
6. Code for WCGNA
7. Additional WCGNA plots

a. Rob and Gos module-trait correlations
b. Rob and Gos module similarity
c. Combined cluster dendrogram
d. Combined gene-trait relationships
e. Rob and Gos gene-trait relationships

## Acknowledgements

This work was supported by the Howard Hughes Medical Institute (DIB). We thank John Lovell and Marie Strader for helpful comments during data analysis and writing.

